# Skin conductance rise time and amplitude discern between different degrees of emotional arousal induced by affective pictures presented on a computer screen

**DOI:** 10.1101/2020.05.12.090829

**Authors:** Miroslava Jindrová, Martin Kocourek, Petr Telenský

## Abstract

Skin Conductance Response (SCR) is a phasic change in electric conductivity of the skin, occurring either non-specifically, or in response to a stimulus (event-related, or ER-SCR). It has long been understood that ER-SCR amplitudes are greater when associated with unpleasant or high-arousal stimuli; however, the relationship between emotional valence and arousal to other ER-SCR measures such as ER-SCR latency (interval between stimulus onset and ER-SCR onset) and ER-SCR rise time (interval between ER-SCR onset and peak amplitude) is less well-established. Here, we presented 60 emotive pictures from IAPS and NAPS affective picture systems to a group of 100 young, healthy adults (50 male and 50 female) and recorded their electrodermal activity. We found that higher emotional arousal was associated with greater ER-SCR amplitudes and shorter ER-SCR rise times. Interestingly, while the increase in ER-SCR amplitudes was only observed for a subset of high-arousal stimuli (score 7-9 on a 9 point scale), the effect on ER-SCR rise times was more graded and particularly sensitive to the difference between low (score 1-3) and medium-arousal (score 4-6) categories. Next, we found that while ER-SCR amplitudes were greater in response to unpleasant stimuli (valence score 1-3 on a 9-point scale), none of the ER-SCR measures could distinguish between neutral (score 4-6) and positive stimuli (score 7-9). We suggest that the increase in ER-SCR amplitudes for unpleasant stimuli is primarily driven by the inherent association between unpleasantness and high arousal. In conclusion, we demonstrate that ER-SCR rise time conveys valuable information about emotional arousal and represents a useful complementary measure to ER-SCR amplitude in order to discern between multiple degrees of emotional arousal. Furthermore, this study confirms the cross-cultural validity of the IAPS and NAPS databases in a sample of young adult Czechs.

## Introduction

Emotions are central to human decision-making and emotional disbalances are commonly associated with neuropsychiatric conditions including depression, anxiety disorders and neurodevelopmental and neurodegenerative disorders. The study of human emotions is thus applicable to a wide array of fields, including neuropsychiatry, clinical psychology, and neuropharmacology but also to areas like neuroeconomics, affective computing or marketing research. Traditionally, research into human emotion relies on the use of verbal self-reporting techniques such as questionnaires and interviews, which however bring several limitations. These include various cultural, age-related, and education-dependent differences in emotional processing and the ability to communicate affective states (Mesquita and Frijda, 1992; Kitayama, 2005; Grühn and Scheibe, 2008; Demenescu et al., 2014; Simoës-Perlant et al., 2018). Although some limitations can be ameliorated by the use of visual self-assessment scales such as the Self-Assessment Manikins (SAMs; Lang, 1980; Bradley and Young, 1994), other issues will persist, such as inaccurate recall of emotional states (Levine and Safer, 2002), deliberate manipulation of self-assessment to avoid shame (Kassam and Mendes, 2013), or the fact that labeling of emotions may impact subsequent emotional experience (Kircanski et al., 2012; Matejka et al., 2013; Torre and Lieberman, 2018). One way to address these limitations is to study psychophysiological responses to emotive stimuli associated with affective processing either at the level of central (e.g. fMRI, EEG) or peripheral nervous systems (e.g. HRV, EDA).

While neuroimaging techniques such as functional magnetic resonance imaging (fMRI) or magnetoencephalography (MEG) represent the state-of-the-art in psychophysiology and provide high-information content data, these methods require expensive medical equipment and specialized expertise and are thus extremely low-throughput, cost-prohibitive, non-transportable and often necessitate grueling logistics. Another limitation of fMRI is the necessity of laying the subject down in an unnatural, noisy and potentially claustrophobic environment, which can influence emotional experience, albeit majority of fMRI subjects report the experience to be tolerable (Szameitat et al., 2009; Hadidi et al., 2014). An alternative approach is to measure peripheral biosignals modulated by autonomous nervous system activity, usually cardiac or electrodermal responses. While the information content is significantly lower, it represents a low-cost, portable, and high-throughput alternative.

The phenomenon of electrodermal response, whereby electric properties of the human skin change as an effect of emotional stimulation, has been known since the late 19^th^ century (Féré, 1888). It is considered to be a result of the activity of eccrine sweat glands which have both cholinergic and adrendergic innervation (Sokolov et al., 1980; Shields et al., 1987). Today, two main components of electrodermal activity are commonly studied; a slower tonic component known as the Skin Conductance Level (SCL) and a much faster phasic component called Skin Conductance Response (SCR; Fowles et al., 1981). The Skin Conductance Response can be characterized by multiple measures, which include SCR amplitude (peak increase in conductance), SCR latency (time interval between stimulus delivery and SCR onset), and SCR rise time (time interval between SCR onset and peak amplitude; Fig. 1B). Highly arousing or unpleasant stimuli are generally understood to be associated with increased SCR amplitudes and shorter latencies and rise times (Dawson et al., 2007). While increased SCR amplitude is the most frequently used SCR correlate of emotional arousal or unpleasantness of stimuli, the specific information value of SCR rise time is less well-understood, although some authors argued for its importance due to its association with attention and cardiac activity already several decades ago (Venables et al., 1980; Shibagaki and Yamanaka, 1991). SCR latency is most frequently used solely as a criterion of SCR specificity to presented stimulus. Current consensus suggests that SCR should only be considered specific (or event-related, hence ER-SCR) if SCR latency lies within a 1-4s time interval (Boucsein et al., 2012). However, recent study suggests that apart from its usefulness as a specificity criterion, SCR latencies vary across individuals and gender groups and may be modulated by associative memory processing (Sjouwerman and Lonsdorf, 2019).

**Fig. 1.**
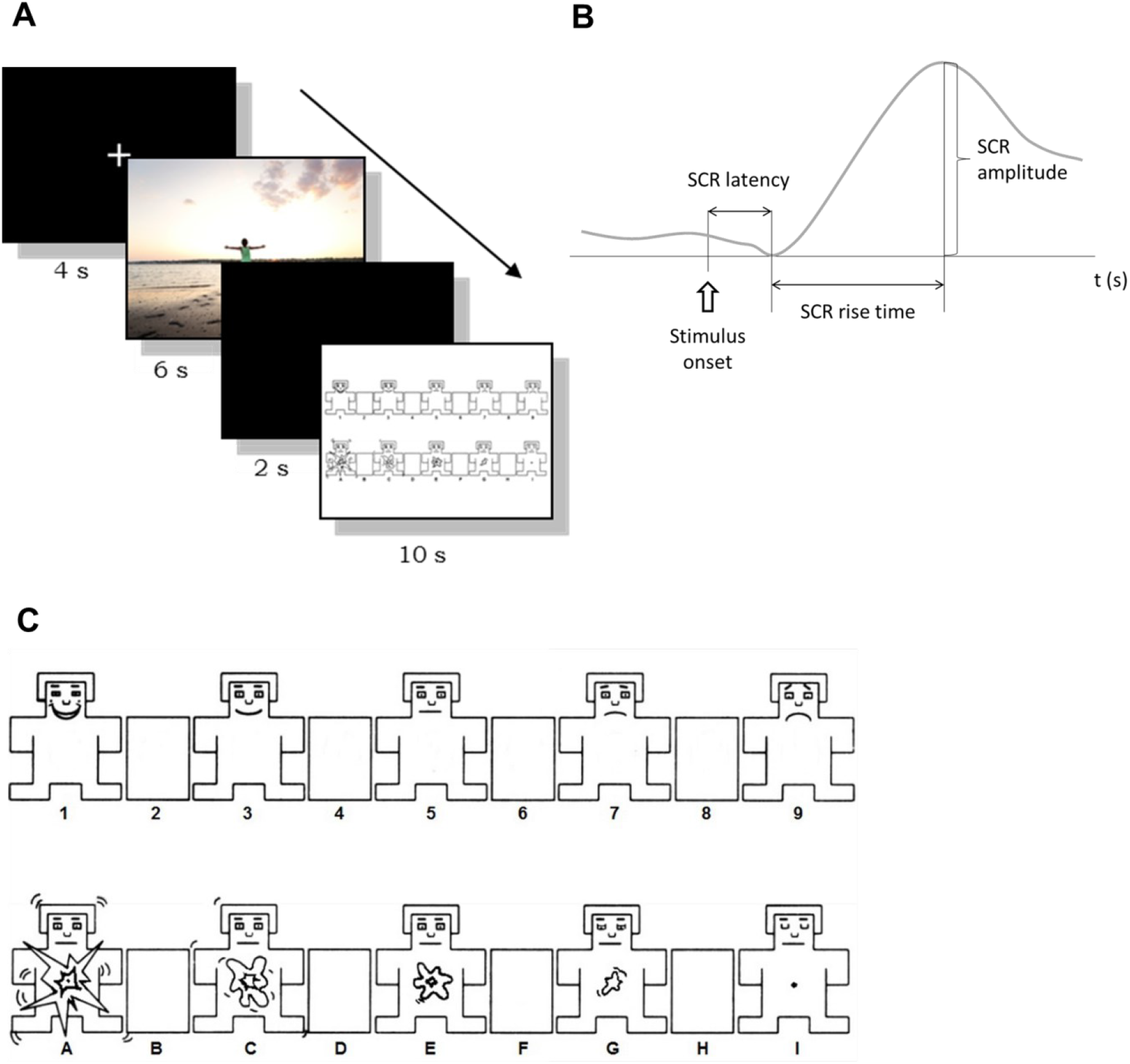
Experimental design of the study. **A:** Timeline of stimulus presentation. Each stimulus was preceded with black screen with a fixation cross (4 s). Subsequently, emotive picture was presented for 6 s, followed by a black screen for 2 s. Last, subjects were asked to subjectively evaluate the emotional valence and arousal of the image, while self-assessment manikins (SAM) were presented on a screen for 10 s. To avoid confusion, valence SAMs were labeled by numbers (1-9), while arousal SAMs were labeled with letters (A-I) and subjects were asked to read aloud the number and letter adjacent to their selected SAMs. **B:** Schematic visualization of the evaluated SCR parameters (redrawn from Dawson et al., 2001). SCR latency is the period between stimulus onset and the onset of SCR. SCR rise time is the period between the SCR onset and the time of max. SCR amplitude. **C:** Detailed view of the self-assessment manikin (SAM) screen used in the experiment. Valence and arousal manikins are in top and bottom rows, respectively.

In the present study, we aimed to investigate whether, in addition to ER-SCR amplitude, ER-SCR latencies and rise times can provide further discriminatory power to distinguish ER-SCRs along categories of emotional valence and arousal. To this end, we presented a total of 60 affective pictures from Nencki Affective Picture System (NAPS; Marchewka et al., 2013) and International Affective Picture Systems (IAPS, Lang et al., 1999) to a group of 100 young (age 18-29), healthy volunteers, recruited in Prague, Czech Republic. We then observed the relationships between ER-SCR measures (amplitude, latency, and rise time) and three categories of emotional valence (unpleasant or negative valence, neutral, and pleasant or positive valence) and arousal (low, medium, high). Included in our analysis were both the standardized ratings provided from NAPS and IAPS databases as well as subjective self-assessment by our subjects. In accordance with previous studies, we found that higher emotional arousal was associated with greater ER-SCR amplitudes and shorter ER-SCR rise times. However, the former measure responded specifically to high arousal stimuli while the latter was more gradual and responded to the difference between low and medium arousal categories. While ER-SCR amplitudes alone could only discern between high-arousal and remaining stimuli, all three categories could be visually represented and statistically distinguished by plotting normalized SCR amplitudes against SCR rise times. Next, we found that while ER-SCR amplitudes were greater in response to negative-valence (unpleasant) stimuli, none of the ER-SCR measures could distinguish between neutral and positive stimuli. We suggest that this observation might be solely explained by the association between unpleasantness and high emotional arousal, which we demonstrate in both our behavioral data and the standardized NAPS and IAPS ratings.

Taken together, this study shows that ER-SCR rise times convey valuable information about emotional arousal and in conjunction with ER-SCR amplitudes allow for better discrimination of skin conductance responses to stimuli with varying degrees of emotional arousal.

## Materials and methods

### Participants and procedure

One hundred healthy participants (50 males, 50 females), aged 18-29 (M=22.82, SD = 2.27) participated in this study. Excluding criteria were any past or present psychiatric or neurological illnesses. Subjects were asked to refrain from consumption of alcohol, caffeine and nicotine on the day of the experiment. Eleven participants were left-handed and all participants had normal or corrected-to-normal vision. Participants signed an informed consent about the participation and were paid for their participation. The experimental sessions were conducted with each participant separately. At the beginning of the session, participants were informed about the experiment, their questions were answered and the informed consent was signed. Participants were then prepared for psychophysiological measurements and given instructions for the testing blocks. The experiments were approved by the Institutional Review Board of the Faculty of Science, Charles University.

### Stimulus presentation

Stimuli were presented on a 48×27 cm screen using eye-tracking software Tobii Studio and observation of presented stimuli was verified with remote eye-tracker (Tobii X2-30) mounted on the bottom screen bezel. Participants were seated at the distance of approx. 60 cm from the screen. Presentation of emotive stimuli comprised of 30 IAPS (International Affective Picture System) and 30 NAPS (The Nencki Affective Picture System) colored pictures. The presented stimuli were chosen so that they uniformly covered a whole range of standardized arousal and valence ratings (Tab. 1, Fig. 2). Before each emotional stimulus, a white fixation cross on a black background was presented for 4 s. Subsequently, the stimulus was presented for 6 s followed by a black screen for 2 s and a screen with valence and arousal SAM (Self-Assessment manikin) scales (Bradley and Lang 1994) for 10 seconds (Fig. 1A and C). Participants were instructed to rate their subjective emotion they experienced while watching the picture and to give their ratings by reading out loud by a number for valence and a letter for arousal (see Fig. 1C). The ratings were marked down in by the experimenter. The instructions for rating were taken from the IAPS protocol and translated into Czech.

**Table 1.**
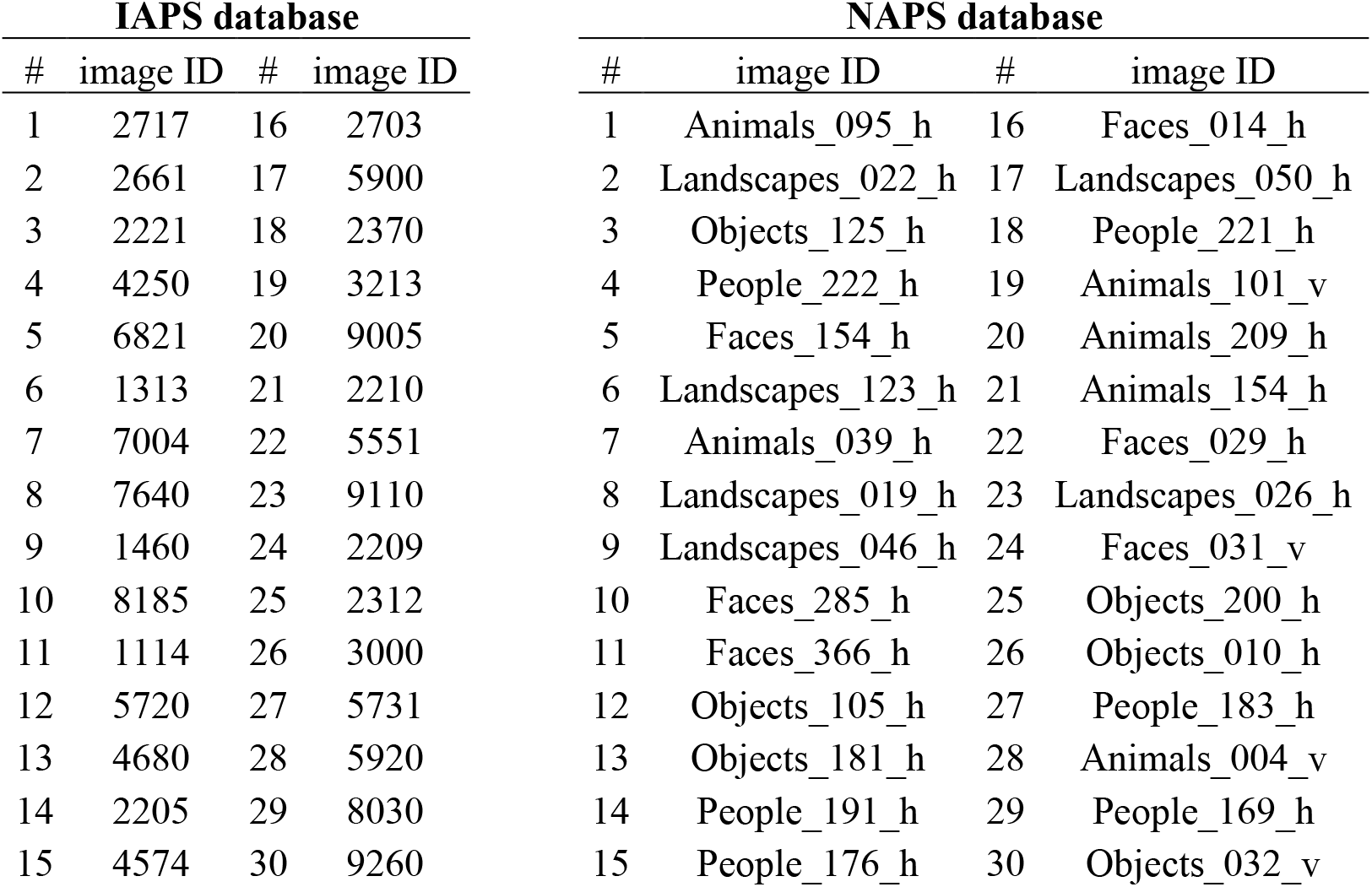
List of selected stimuli from the IAPS and NAPS databases. 30 stimuli from each database were selected. Image number (#) denotes order in which the pictures were presented within the IAPS or NAPS block.

| IAPS database |  |  |  | NAPS database |  |  |  |
| --- | --- | --- | --- | --- | --- | --- | --- |
| # | image ID | # | image ID | # | image ID | # | image ID |
| 1 | 2717 | 16 | 2703 | 1 | Animals_095_h | 16 | Faces_014_h |
| 2 | 2661 | 17 | 5900 | 2 | Landscapes_022_h | 17 | Landscapes_050_h |
| 3 | 2221 | 18 | 2370 | 3 | Objects_125_h | 18 | People_221_h |
| 4 | 4250 | 19 | 3213 | 4 | People_222_h | 19 | Animals_101_v |
| 5 | 6821 | 20 | 9005 | 5 | Faces_154_h | 20 | Animals_209_h |
| 6 | 1313 | 21 | 2210 | 6 | Landscapes_123_h | 21 | Animals_154_h |
| 7 | 7004 | 22 | 5551 | 7 | Animals_039_h | 22 | Faces_029_h |
| 8 | 7640 | 23 | 9110 | 8 | Landscapes_019_h | 23 | Landscapes_026_h |
| 9 | 1460 | 24 | 2209 | 9 | Landscapes_046_h | 24 | Faces_031_v |
| 10 | 8185 | 25 | 2312 | 10 | Faces_285_h | 25 | Objects_200_h |
| 11 | 1114 | 26 | 3000 | 11 | Faces_366_h | 26 | Objects_010_h |
| 12 | 5720 | 27 | 5731 | 12 | Objects_105_h | 27 | People_183_h |
| 13 | 4680 | 28 | 5920 | 13 | Objects_181_h | 28 | Animals_004_v |
| 14 | 2205 | 29 | 8030 | 14 | People_191_h | 29 | People_169_h |
| 15 | 4574 | 30 | 9260 | 15 | People_176_h | 30 | Objects_032_v |

**Fig. 2.**
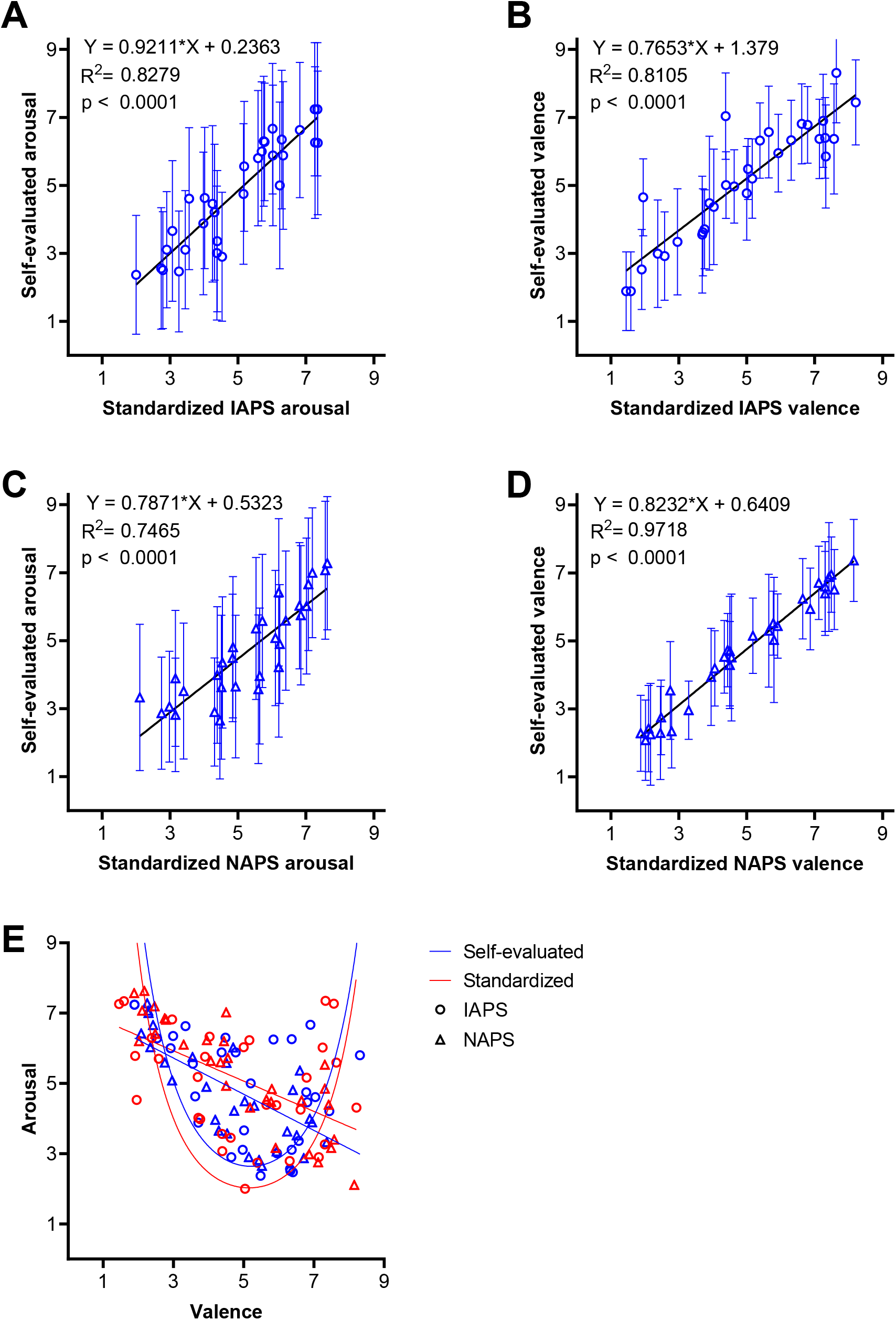
Validation of the emotive picture sets. **A-D:** Mean subjective ratings of valence and arousal were significantly correlated to the standardized valence and arousal mean values obtained from IAPS and NAPS databases. Error bars represent standard deviation. Parameters of linear regression are shown. **A:** Linear relationship between standardized and self-evaluated arousal values in the IAPS subset. **B:** Relationship between standardized and self-evaluated valence values in the IAPS subset. **C:** Relationship between standardized and self-evaluated arousal values in the NAPS subset. **D:** Relationship between standardized and self-evaluated valence values in the NAPS subset. **E:** Summary plot of all affective pictures from the IAPS (circles) and NAPS (triangles) databases plotted as mean values of valence and arousal. Standardized values taken from the respective databases are shown in red, self-evaluated ratings are shown in blue. The curves and lines in the corresponding colors represent quadratic and linear association between the emotional dimensions, respectively.

### Electrodermal activity recording and data analysis

Electrodermal activity was recorded with disc adhesive passive electrodes (Medimaxtech) placed on the palmar side of the medial phalanges of the index and middle finger of the non-dominant hand. Both electrophysiological measures were recorded at a sampling rate 5000 Hz using Neuron-Spectrum 4/P acquisition system (Neurosoft). We then evaluated SCR amplitudes, latencies and rise times. Only SCRs with latencies within 1-4 s and peak SCR amplitudes > 0.01 μS were included in the analysis as ER-SCRs. Stimulus presentation was synchronized with data acquisition using StimTracker (Cedrus Corporation, San Pedro, CA, USA).

Three participants had to be excluded from analysis due to incomplete or low-quality recordings. Electrodermal data from the remaining 97 participants were included in evaluation and statistical analysis. Raw electrodermal data were first exported for analysis to Acqknowledge 5.0 software (Biopac systems Inc., Goleta, CA, USA). SCR data were then exported to Microsoft Excel. To reduce inter-individual variability, SCR amplitudes were normalized by converting to z-scores according to the following formula:

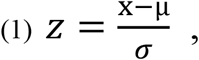

where z is the normalized score, x is the observed amplitude, μ is the mean of all ER-SCR amplitudes included in analysis from the test subject, σ is the standard deviation of the ER-SCR amplitudes from the subject. Final statistical analysis was performed in GraphPad Prism 8 (GraphPad Software, San Diego, CA, USA).

To analyze the ER-SCR measures in relation to emotion, we first categorized the 9-point self-assessment scale scores into 3 subcategories; low (scores 1-3), medium (4-6) and high (7-9) for arousal and negative/unpleasant (1-3), neutral (4-6) and positive/pleasant (7-9) for valence. The standardized average ratings from NAPS and IAPS databases were categorized using range intervals: valence scores (V) 1≤V≤3.67 were categorized as negative valence, 3.67<V≤6.33 as neutral, and 6.33<V≤9 as positive valence category. Standardized arousal scores were analogously grouped into low, medium and high arousal categories.

Ordinary one-way ANOVA and Tukey’s multiple-comparison *post-hoc* tests were used to determine the effects of arousal and valence on ER-SCR characteristics. Linear regression analyses were used to determine the correlation between subjective self-assessments and standardized valence and arousal ratings. Linear and quadratic regressions were used to plot the linear and quadratic relationship between valence and arousal (Fig. 2E).

## Results

### Cross-cultural validity of NAPS and IAPS stimuli

We first performed linear regression to determine whether the standardized NAPS and IAPS ratings correlated with the self-assessment scores by our subjects (Fig. 2A-D). For both NAPS and IAPS stimuli, self-assessment valence and arousal scores were highly correlated with standardized ratings (p<0.0001, R^2^ = 0.75-0.97).

### Effect of emotive arousal on electrodermal responses

We then investigated the effect of arousal on ER-SCR amplitudes, latencies and rise times (Fig. 3). When categorized according to self-assessment scores, arousal significantly influenced all 3 studied ER-SCR measures (Fig. 3A). One-way ANOVA found significant main effects of arousal on z-score normalized ER-SCR amplitude (F (2, 1642) = 10.81; p<0.0001), ER-SCR latency (F (2, 1643) = 7.993; p<0.0001) and rise time (F (2, 1643) = 7.051; p<0.0001). Tukey’s multiple comparison *post hoc test* revealed significant differences between high and low and between high and medium arousal categories (both, p<0.001). In ER-SCR latency, *post hoc* test showed differences between high and low (p<0.05) and high and medium (p<0.001) categories. For ER-SCR rise times, we found a difference between low and medium (p<0.05) and low and high (p<0.001) categories. As to the standardized ratings-based categories, significant main effect of arousal was found in ER-SCR amplitudes (F (2, 1642) = 5.452; p<0.01) and rise time (F (2, 1643) = 2.809; p<0.05); *post hoc* tests revealed significant difference between low and medium arousal (p<0.05) and low and high arousal in ER-SCR amplitudes, and between high and low arousal in SCR rise times (Fig. 3B).

**Fig. 3.**
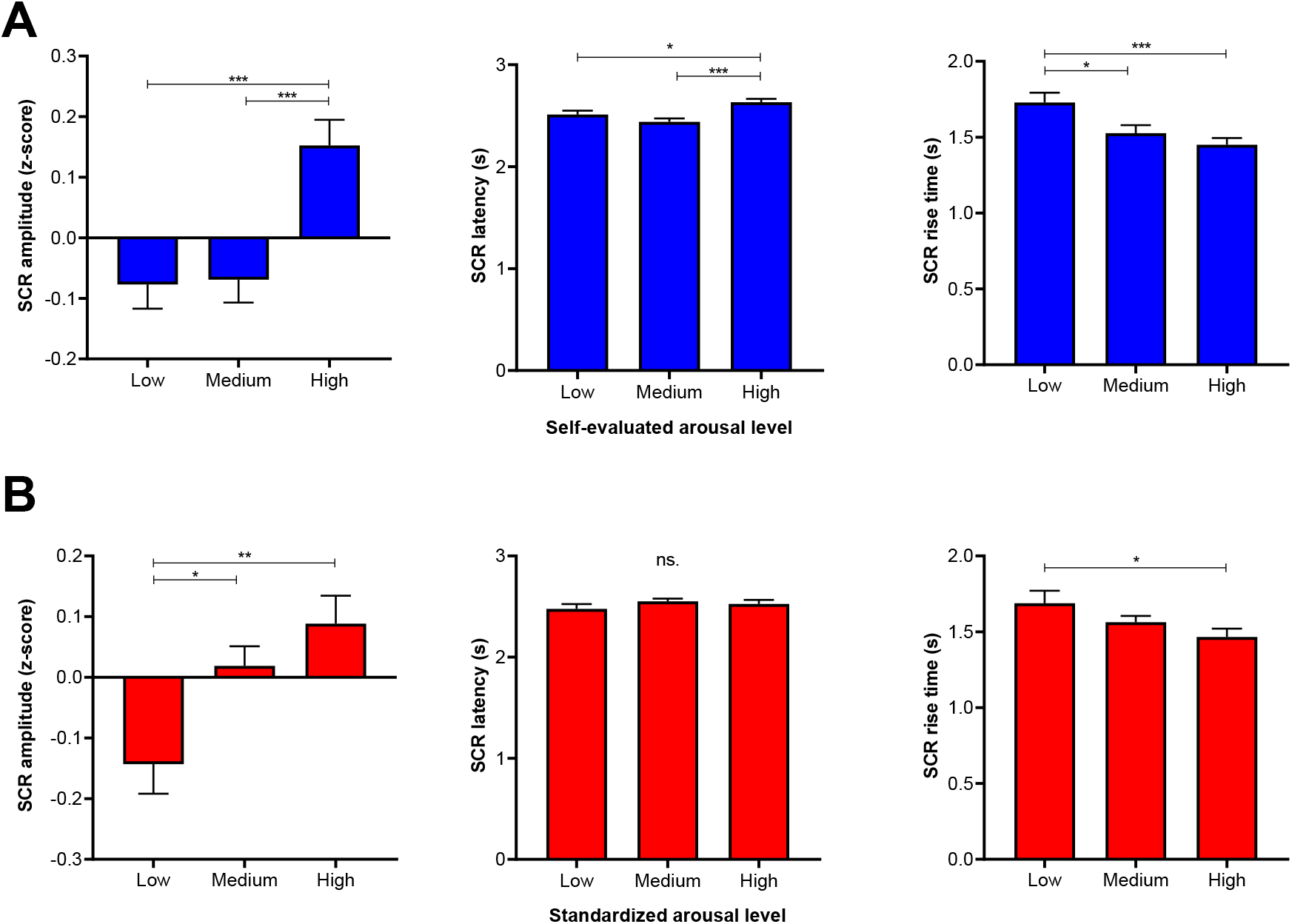
Properties of skin conductance responses to low, medium and high arousal stimuli. **A:** Categories of emotional arousal represented as self-evaluation by subjects -(low = score 1-3, medium = score 4-6, high = score 7-9). **B:** Categories of emotional arousal represented as standardized average ratings from IAPS and NAPS databases. Following intervals of average ratings were used for each category: 1≤x≤3.67 for low, 3.67<x≤6.33 for medium, and 6.33<x≤9 for high arousal category. (* = p < 0.05; ** = p < 0.01; *** = p < 0.001; ns. = non-significant)

### Effects of emotive valence on electrodermal responses

Subsequently, we investigated whether ER-SCR amplitudes, latencies and rise times varied across valence categories for both self-assessment scores and standardized IAPS and NAPS ratings (Fig. 4). One-way ANOVA showed significant main effect of self-assessed valence on normalized ER-SCR amplitudes (F (2, 1641) = 3.431; p<0.05). Tukey’s multiple comparison *post hoc* test revealed significant difference (p<0.05) between negative and neutral categories. We found no differences in ER-SCR latencies and rise times between the self-assessed valence categories. Interestingly, more significant differences were found across the valence categories based on standardized scores. One-way ANOVA showed significant main effect of valence on ER-SCR amplitudes (F (2, 1642) = 5.066, p<0.01) and Tukey’s multiple comparison *post hoc* test showed significant differences between negative and neutral and negative and positive categories (both p<0.05). Furthermore, we found significant main effect of valence on ER-SCR latency (F (2, 1643) = 3.193; p<0.05) and *post hoc* test revealed differences between negative and positive category.

**Fig. 4.**
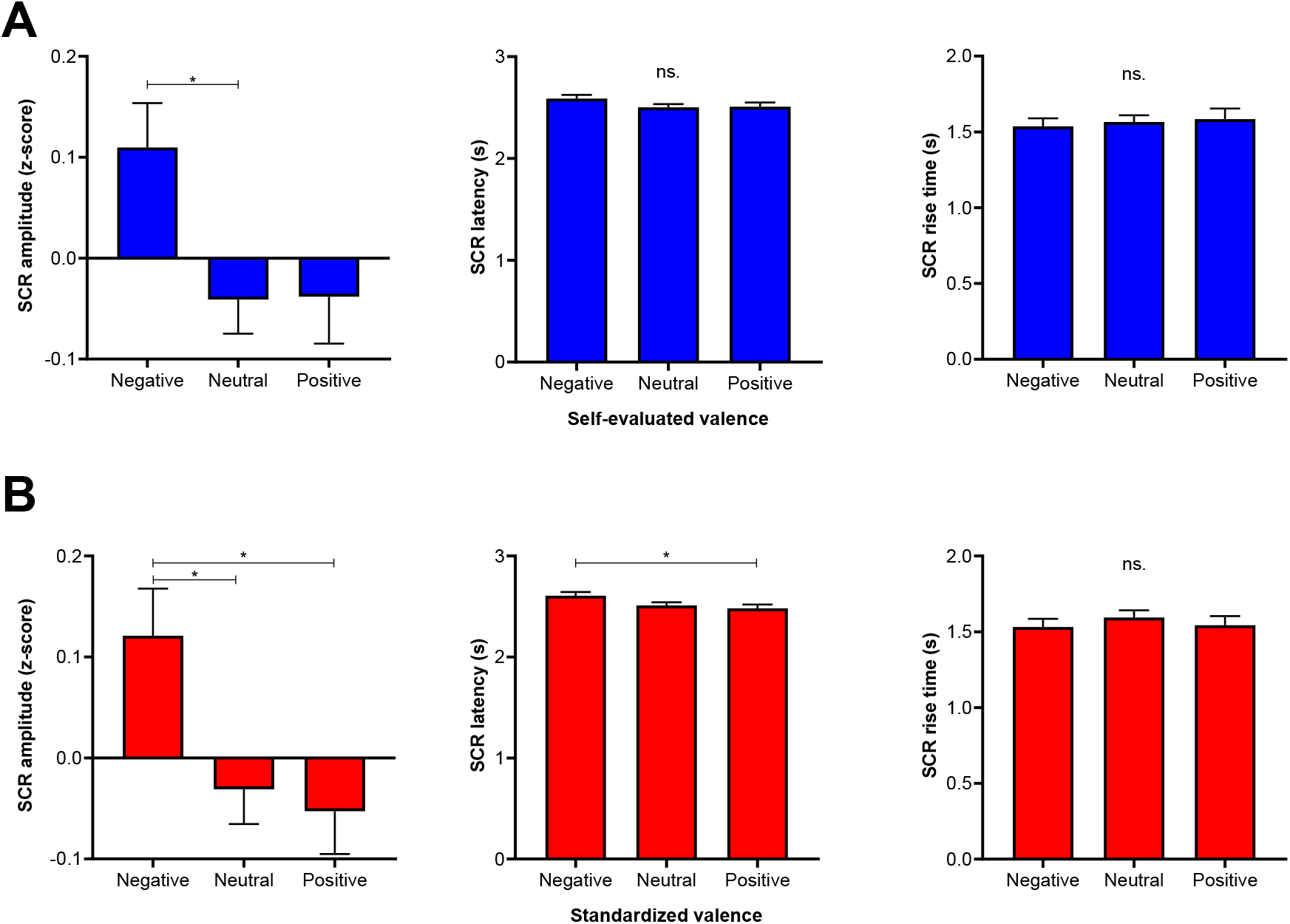
Properties of skin conductance responses to stimuli with negative, neutral and positive emotional valence. **A:** Categories of emotional valence represented as self-evaluation by subjects -(negative = valence 1-3, neutral = valence 4-6, positive = valence 7-9). **B:** Categories of emotional valence represented as standardized average ratings from IAPS and NAPS databases. Following intervals of average ratings were used for each category: 1≤x≤3.67 for negative, 3.67<x≤6.33 for neutral, and 6.33<x≤9 for positive valence category. (*= p < 0.05; ns. = non-significant)

## Discussion

This work has demonstrated that ER-SCR rise time conveys valuable information about emotional arousal and represents a useful complementary measure to ER-SCR amplitude in order to discern between multiple degrees of emotional arousal.

Since this work is to our knowledge one of a few studies to use the IAPS (Czékoová et al., 2015) and the very first to use the NAPS affective picture databases in the Czech cultural context, we first investigated whether there is a good correlation between self-assessment scores of valence and arousal as collected from our subjects during the experimental session and the standardized valence and arousal ratings, which are available in the IAPS and NAPS databases. Although a full validation of the stimulus sets was far beyond the scope of this work, our correlations indicate a high cross-cultural validity of NAPS and IAPS stimuli.

We subsequently used both the self-assessed scores and standardized ratings to categorize stimuli into negative, neutral and positive categories of valence, and low, medium and high categories of arousal. As expected, the self-assessment arousal categories led to more significant differences and affected all 3 of the studied ER-SCR measures. Importantly, we found that while the SCR amplitude increase was only detectable in the high-arousal category, the effect on SCR rise times was not only more gradual, but also particularly sensitive to a difference between low and medium categories. Therefore, a combination of ER-SCR rise time and amplitude could be used to effectively discern between the 3 categories of emotional arousal, where the former measure can distinguish between low and medium arousal and the latter between medium and high arousal levels. This can be graphically illustrated with a two-dimensional Cartesian plot of ER-SCR rise time against normalized ER-SCR amplitude (Fig. 5A). Increasing emotional arousal follows an “L”-shaped trajectory: a decrease in rise time along the y-axis is followed by an increase in ER-SCR amplitude along the x-axis. Although, to our knowledge, this effect wasn’t previously discussed or directly reported, a recent study by Bari and colleagues (2018) shows similar results. The researchers used 3 levels of electric nociceptive stimuli and evaluated ER-SCR responses. While rise times were significantly changed between stimulus intensities 1 and 2, amplitudes changed between intensities 2 and 3. This is not only in accord with our observations, but it may also suggest that this phenomenon could be applicable across stimulus modalities; however, further studies are needed to support this hypothesis. The above described phenomenon could be attributable to the influence of attention. It is reasonable to assume that stimuli eliciting medium levels of arousal are commanding increased levels of attention compared to low-arousal stimuli. Previous studies indeed reported that rise times are inversely associated with attention and attentional openness and positively associated with defensive or closed attentional stance (Venables et al, 1980; Shibagaki and Yamanaka, 1991).

**Fig. 5.**
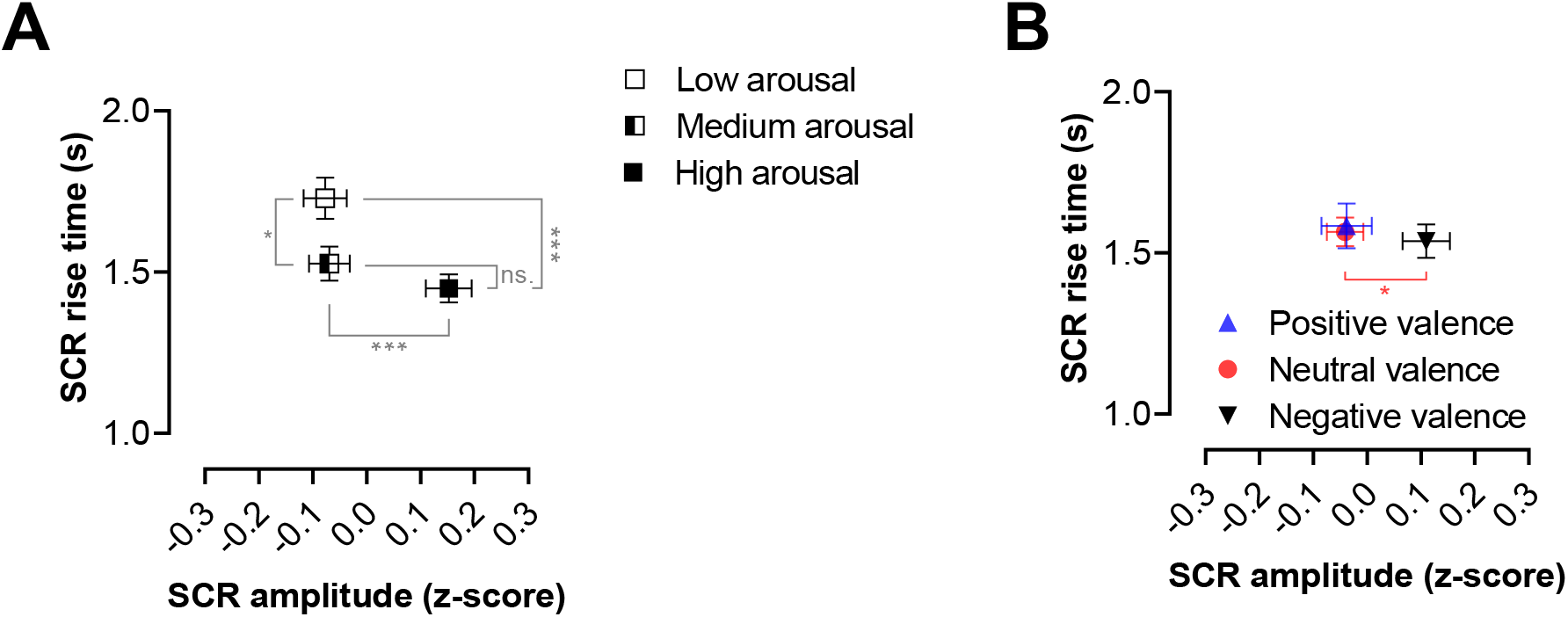
Two-dimensional plots of normalized SCR amplitude and rise time for categories of emotional valence and arousal. All data and statistical test results indicated in this figure are identical to those reported in figures 3 and 4. **A:** All 3 arousal categories can be differentiated by two dimensional analysis (SCR rise time x normalized SCR amplitude). Increasing emotional arousal follows approximately the shape of the letter “L”: SCR rise times are particularly sensitive to the difference between low and medium arousal categories, while SCR amplitude was increased in response high-arousal stimuli. **B:** Images of positive and neutral valence could not be differentiated by SCR properties. Only negative-valence pictures were associated with increased normalized SCR amplitudes, which could possibly be due to the interdependence of negative valence and high emotional arousal. Error bars represent SEM. (* = p < 0.05; *** = p < 0.001; ns. = non-significant)

We further studied how ER-SCR parameters responded to categories of emotional valence. Within categories of self-evaluated valence, we only found an effect on ER-SCR amplitudes, which were significantly increased in the negative valence category. This is in accordance to a generally accepted view that increased ER-SCR amplitude is associated with unpleasant stimuli (Dawson et al., 2007). No other effects of self-assessed valence on ER-SCRs were seen, which we also illustrated by an amplitude/rise time plot (Fig. 5B). Negative valence is associated with (and cannot be separated from) high levels of arousal, which we also demonstrated in our sample (Fig. 2E). Thus, we can argue that the effect of valence on ER-SCR is mediated by the relation between negative valence and arousal. This is in accord with the observation, that SCR amplitude is associated with amygdalar activation elicited by threatening stimuli (Wood et al., 2014). Interestingly, we found more significant differences in ER-SCR parameters for valence categories derived from standardized NAPS and IAPS ratings. However, these very differences could be observed as ‘non-significant trends’ among the self-assessed categories as well. Hence, we reason that this is due to a very good agreement between self-assessed and standardized ratings of valence. Standardized valence ratings of NAPS stimuli correlated with average self-assessed scores particularly well (R^2^=0.9718; Fig. 1D). Since the NAPS database originated in Poland, a possible interpretation is that this might be due to the relative cultural similarity between the neighboring Poles and Czechs.

In conclusion, our study shows that ER-SCR rise time, a parameter often omitted from reports of electrodermal activity, has an excellent potential as a marker of low to medium levels of emotional arousal. We argue that our work necessitates the incorporation of ER-SCR rise time in further studies of phasic electrodermal responses.

## Acknowledgements

This study was supported by GAUK grant 1204217, Junior GAČR grant no. 16-21228Y, PRIMUS/SCI/33 and by the European Regional Development Fund - Project ENOCH (No. CZ.02.1.01/0.0/0.0/16_019/0000868). The authors declare no conflict of interests.

